# Spatio-temporal modeling reveals a layer of tunable control circuits for the distribution of cytokines in tissues

**DOI:** 10.1101/2022.03.17.484722

**Authors:** Patrick Brunner, Lukas Kiwitz, Kevin Thurley

## Abstract

Cytokines are diffusible mediators of cell-cell communication among immune cells with critical regulatory functions for cell differentiation and proliferation. Previous studies have revealed considerable spatial inhomogeneities in the distribution of cytokine molecules in tissues, potentially shaping the efficacy and range of paracrine cytokine signals. How such cytokine gradients emerge and are controlled within cell populations is incompletely understood. In this work, we employed a spatial reaction-diffusion model to systematically investigate the formation and influence of spatial cytokine gradients. We found the fraction of cytokine secreting cells to be the main source of spatial inhomogeneity and subsequent activation. Positive feedback from local cytokine levels upon cytokine receptor expression leads to further increased spatial cytokine inhomogeneities. By exploring the effect of co-clustering cytokine secreting cells and cells with large amounts of receptor expression, as in the presence of regulatory T cells in the vicinity of antigen-presenting cells, we found that such constrained tissue architecture can have profound effects on the range of paracrine cytokine signals.

## Introduction

Cytokines shape the differentiation decisions of T helper (T_h_) cells into subtypes. While bulk measurements of cytokine concentration (1) and cellular response to cytokine signals (2,3) are experimentally accessible, the spatial *in vivo* cytokine distribution has proven challenging to probe in vivo. A common assumption is that cytokine concentrations equilibrate quickly across the typical spatial dimensions of lymph nodes or other lymphoid tissue compartments, since cytokines are small and highly diffusible proteins. However, we found in previous model simulations that substantial spatial inhomogeneities in cytokine concentration can occur in a physiologic parameter regime (Table 1), and that such inhomogeneities can induce a substantial increase in the efficacy of paracrine cytokine signaling (4,5). These results are in agreement with recent experimental studies indicating substantial and tunable cytokine inhomogeneities in lymphoid tissue (6–9). One reason for spatial cytokine inhomogeneities despite fast diffusion is that the time required for a cytokine molecule to find an unoccupied receptor is much shorter than the diffusion time over distances on the tissue scale (5). Hence, if cytokine secretion is roughly balanced with the uptake ability of the surrounding cells, stable gradients between cytokine sources and sinks can arise despite fast diffusion, similar to the emergence of stable gradients of the electrical potential in a condenser in electrostatics.

**Table 1.**
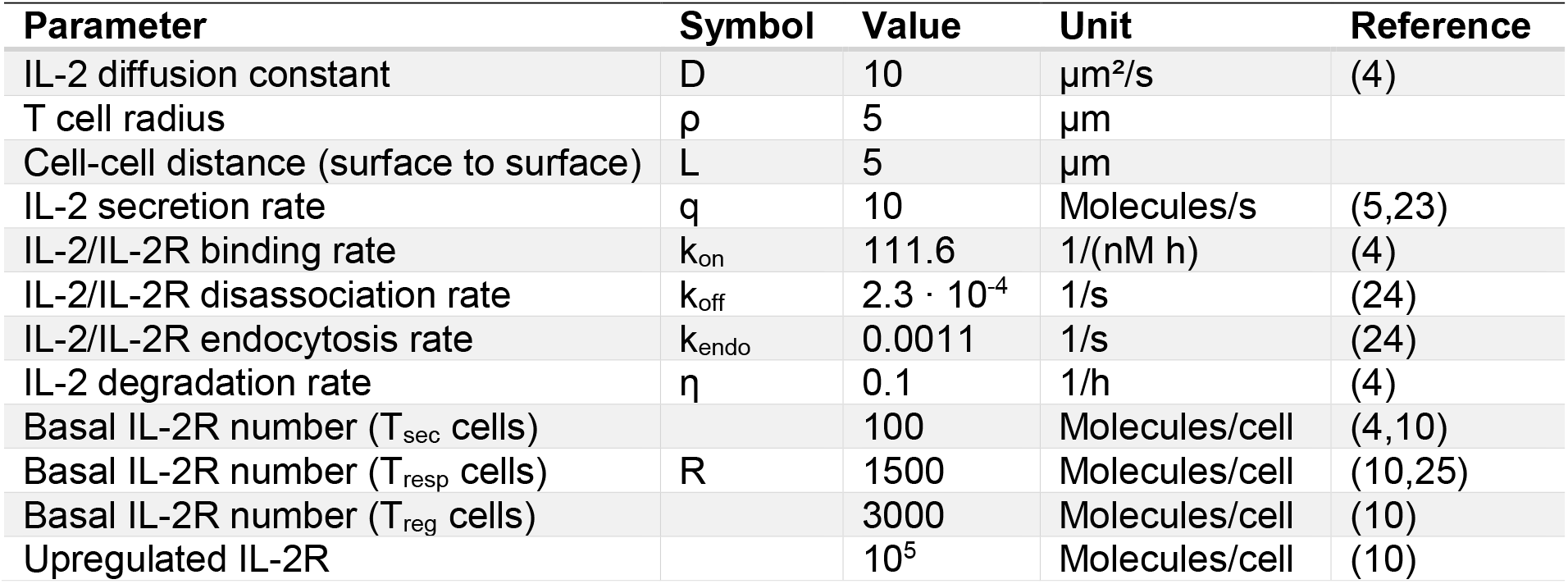
Standard parameter values used in the study.

While those general properties of cytokine diffusion patterns seem well understood, it is less clear which properties of the cell population are critical for the emergence of stable cytokine gradients. In the light of the reasoning above, it seems conceivable that a population of identical cells will not generate cytokine gradients, and rather, local differences in terms of cytokine secretion and uptake capabilities are required. Indeed, data indicate that the strength of T cell receptor engagement is reflected in the number of activated cells rather than the strength of cytokine secretion. As a result, the initial contact with an antigen or the distribution of auto-reactive cells under homeostatic conditions will yield a separation of a T cell population into cytokine secreting cells and cells merely responsive to cytokine signals due to cytokine receptor expression. Another potential source of heterogeneity is the highly diverse amount of receptor expression, which has been found to vary between individual cells by at least 2 orders of magnitude (10).

In addition to such considerations that are relevant for many physiologically relevant scenarios of spatial cell-cell-communication, also dynamic interactions can shape spatial cytokine profiles. In particular, cytokine signaling often imposes feedback on the receptor expression of target cells, such as positive feedback upon signaling through the cytokine interleukin (IL)-2, which leads to increased expression of CD25, the alpha-unit of the high-affinity IL-2 receptor. Consequently, cytokine signaling itself may have an impact on the local cytokine uptake dynamics and therefore on the magnitude of spatial concentration gradients.

Here, to systematically analyze the causes and consequences of stable spatial gradients in cytokine signaling, we derived a general framework that couples a response-time description (11) of cellular signal integration to a spatial description of cytokine diffusion dynamics. We investigated the role of cellular properties such as cytokine secretion rate and heterogeneity of receptor expression, processed to dynamic model simulations including feedback regulation, and finally analyzed the consequences of a dedicated tissue architecture such as co-localization of cytokine secreting cells (T_sec_) and regulatory T cells (T_reg_). We found that the amount of cytokine secreting cells is a major driver of concentration inhomogeneities, which is further amplified by feedback in cell-cell interaction dynamics and subject to fine-tuned regulation by means of cellular localization and uptake dynamics.

## Results

### Spatial inhomogeneities drive cytokine signaling efficacy

In order to systematically quantify spatial cytokine inhomogeneities arising from secretion and uptake dynamics in a randomly distributed cell population, we developed a mathematical model based on reaction-diffusion (RD) mechanics (Figure 1A). Cytokines are produced by a small population of secreting cells, and rapidly diffuse within the extracellular space, where they can bind to cytokine receptors on the surface of responding cells. The cytokine-receptor complex is subsequently internalized, initializing a signaling cascade that results in phosphorylation of a cytokine-specific transcription factor from the STAT family, such as STAT5 in the case of IL-2. Considering the time-scale separation between fast cytokine diffusion and much slower cellular signal processing, the cytokine concentration is evaluated in quasi-steady-state in our simulations.

**Figure 1:**
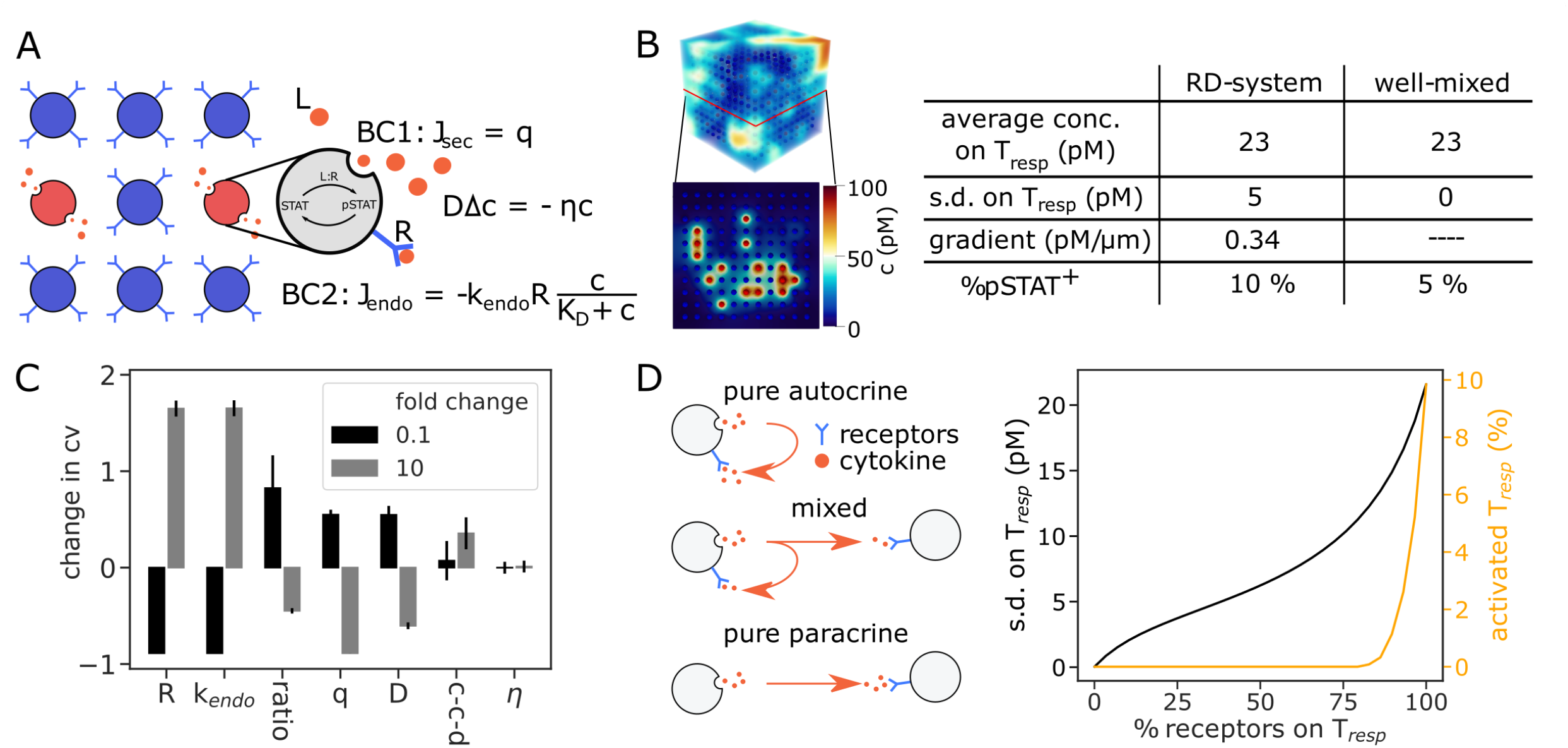
Spatial inhomogeneities induce stable paracrine signaling activity. (A) Schematic of the system setup containing secreting cells (red), responding cells (blue) and cytokines (orange). Cytokines diffuse and decay in the extracellular space while cell-cell-interaction is facilitated via boundary conditions (BC) and pSTAT5 signaling. Cytokines are secreted with rate *q* and internalized through a Michaelis-Menten type endocytosis rate. (B) Left: Setup of the three-dimensional simulation with a cross section showing the secreting cells in red and responding cells in blue. The color bar represents the cytokine concentration in the extracellular space. Right: Comparison of system metrics between the three-dimensional spatial model and a well-stirred model lacking any spatial attributes. “average” and “s.d.”, average and standard deviation across surface concentrations at all responder cells; “gradient”, sum of the gradient over the cytokine field; “%pSTAT+”, percentage of activated responder cells. (C) Scan over system parameters concerning cytokine inhomogeneity, measured by the coefficient of variation (CV) of the surface cytokine concentration across all responder cells. “ratio”, ratio between numbers of secreting and responding cells; “c-c-d”, cell-cell-distance. (D) Scan from pure autocrine (all receptors on secreting cells) to pure paracrine (all receptors on responder cells) signaling, shown is the effect on activation and cytokine inhomogeneity at responder cells.

3D simulations of the model show a spatial pattern containing wells and sinks: the high density of cytokine molecules in the vicinity of secreting cells represents the wells, while the uptake of cytokine by responding cells acts as sinks. As a consequence of the randomized, low-density distribution of cytokine secreting cells, the system shows substantial spatial inhomogeneity in cytokine concentration (Figure 1B). To delineate spatial from non-spatial effects, we compared the RD-system to an ordinary differential equation (ODE) based system, also called well-mixed system since in that case infinitely fast diffusion is assumed so that concentration inhomogeneities disappear. Comparing these two models in terms of cytokine concentration, inhomogeneity and activation, we find that while the average cytokine concentration is preserved, the RD-system doubles the number of activated cells (Figure 1B, right).

To further quantify this effect, we analyzed the change in the coefficient of variation (CV) of the cytokine concentration across the cell surfaces of all responder cells (Figure 1C), as a measure of signaling-effective spatial cytokine inhomogeneity. We identified the cytokine receptor expression *R* and the endocytosis rate *k*_*endo*_, together corresponding to the cytokine uptake capacity of responder cells, as the most important drivers of spatial inhomogeneity, followed by the ratio between secreting and responding cells. Interestingly, other parameters such as the diffusion speed *D* or the cell-cell-distance had a much lower impact on the cytokine concentration inhomogeneity.

An additional consideration in this complex model is the variation of multiple interconnected parameters. Experimental evidence (12) and theoretical considerations (13) suggest the ratio of receptors between secreting and responding cells to determine the mode of cytokine signaling (Figure 1D). If the receptors were all localized on the secreting cells only autocrine signaling is possible, and vice versa. Our model suggests only strong paracrine signaling to enable the activation of surrounding responding cells. This also coincidences with an increase of inhomogeneity in the system, again highlighting its importance.

### All-or-none behavior of cytokine secreting cells as main source of spatial inhomogeneity

To further illuminate the emergence of spatially unequal cytokine distributions in a randomly distributed cell population, we performed a systematic analysis of the major drivers of inhomogeneity: the fraction of secreting cells, the heterogeneity of receptor expression on responding cells and the cytokine uptake dynamics on responding cells (Figure 2A). First, we increased the number of secreting cells in the system while keeping the systemic secretion rate and receptor count constant. That allowed for a constant cytokine concentration, giving us a more accurate representation of the inhomogeneity through the standard deviation. Comparing the models, both showed nearly identical cytokine concentrations over the whole scan. The standard deviation is at its maximum when the cytokine secretion is localized on only one secreting cell and then gradually decreases. Once all cells are cytokine secreting cells, so that all simulated cells are identical (Figure 2B, left), the standard deviation approaches zero as expected.

**Figure 2:**
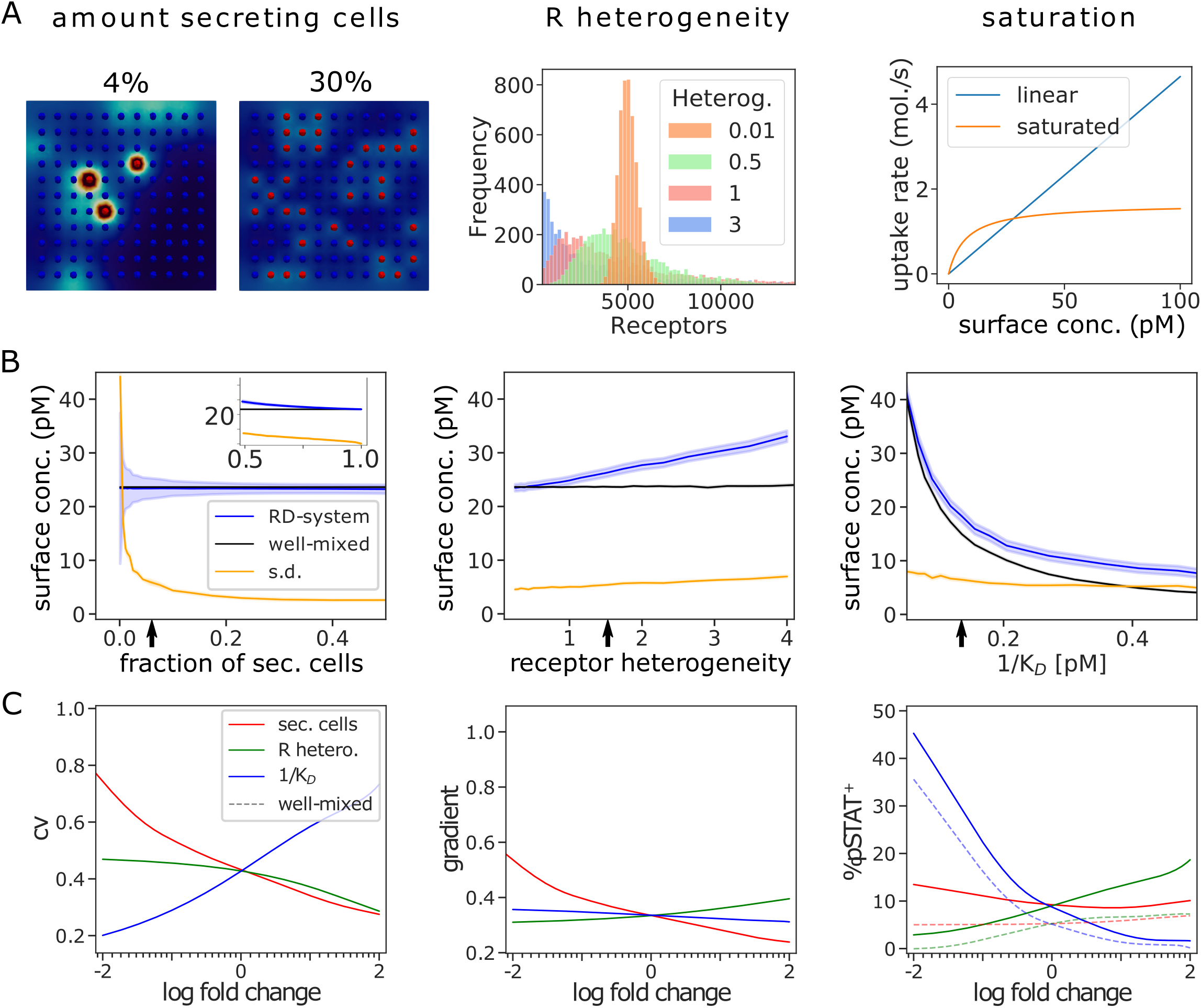
Systematic analysis reveals major drivers of cytokine inhomogeneity. (A) Illustration of the approach. Either the fraction of cytokine secreting cells (left), the amount of receptor heterogeneity (middle) or the degree of saturation in the cytokine uptake rate (right) is varied, while keeping all other parameters constant. (B) Analysis with respect to average and standard deviation of the surface concentration across responder cells. The arrow indicates standard parameter values. (C) Fold-change parameter scan with respect to standard parameter values, indicating the effect on cytokine inhomogeneity (CV and gradient) and paracrine signaling effectivity (%pSTAT+), as in Figure 1B.

To analyze receptor heterogeneity, the receptor count of every cell was log-normally distributed, while varying the standard deviation of the underlying normal distribution and keeping the average receptor count constant (Figure 2A, middle). In the well-mixed model, the localization of receptors does not play any role, so in turn this parameter scan shows no significant effect (Figure 2B, middle). In the RD-system, not only the standard deviation but also the concentration increases with increasing receptor heterogeneity. This seemingly paradoxical increase in concentration can ultimately be attributed to the long-tailed shape of the log-normal distribution, which implies that most cells receive very low and a few cells receive very high receptor counts. In our three-dimensional model these few randomly placed high receptor cells are not sufficient to control the increased cytokine concentrations caused by the overwhelming quantity of low receptor cells, leading to the increase in average cytokine concentration.

Next, we asked how cells respond to different cytokine concentrations by changing from a linear uptake function to a saturated one, while varying the reciprocal of the half saturation constant K_D_ (Figure 2A, right). With increasing 1/K_D_, the cellular cytokine uptake increases, lowering both the average and the standard deviation of the cytokine concentration across cell surfaces on responder cells (Figure 2B, right). Both the RD-system and the well-mixed model show similar behavior in this range, revealing a low influence of the degree of saturation on cytokine inhomogeneity.

Finally, we systematically assessed the effect of the three parameters fraction of secreting cells, receptor heterogeneity, and degree of saturation at cytokine uptake, in the vicinity of our literature-derived set of standard parameters (Figure 2C). To fully capture the spatial cytokine distribution, we evaluated the average gradient across the whole intercellular space in addition to the CV across cell surface. These metrics consistently highlight the fraction of secreting cells to be the main driver for inhomogeneity, followed by K_D_ and receptor heterogeneity (Figure 2C). In terms of activation, 1/K_D_ plays a significant role, which can mainly be attributed to the increase in cytokine concentration associated with its negative fold change. Accordingly, the difference between the RD-system and the well-mixed model remains constant in this regime. For both the receptor heterogeneity and the fraction of secreting cells, an increase in inhomogeneity leads to an increase in activation, indicating again that inhomogeneities facilitate activation.

### Positive feedback on receptor expression modulated spatial cytokine gradients

After analyzing the origin of spatial cytokine gradients in static scenarios, we proceeded to consider the impact of cell-state dynamics on the cytokine concentration field. A scenario that has already evoked considerable interest in the scientific community is positive feedback from cytokine binding to cytokine receptor expression on responder cells, which has been experimentally established for instance for the cytokine IL-2 that signals through pSTAT5 (Figure 3A) (14). In this system, naive T cells (T_naive_) act as the responding cells, while secreting cells correspond to activated naive T cells (here T_sec_). Here, we started with a system of 1000 cells, out of which 5% are T_sec_ (Figure 3B). Considering the pSTAT5 levels of every cell in the system, we found certain cells exhibiting elevated pSTAT5 levels after 2 days, while the remaining cells dropped below our activating threshold of 0.5, consistent with previous work (5) (Figure 3C). That suggests to analyze pSTAT5^+^ and pSTAT5^-^ cells separately (Figure 3D). Due to receptor up-regulation in the pSTAT5^+^ subgroup, the surface concentration of these cells is much lower. Yet, due to the higher amounts of receptors, these diminished concentrations still suffice for stable pSTAT5 up-regulation.

**Figure 3:**
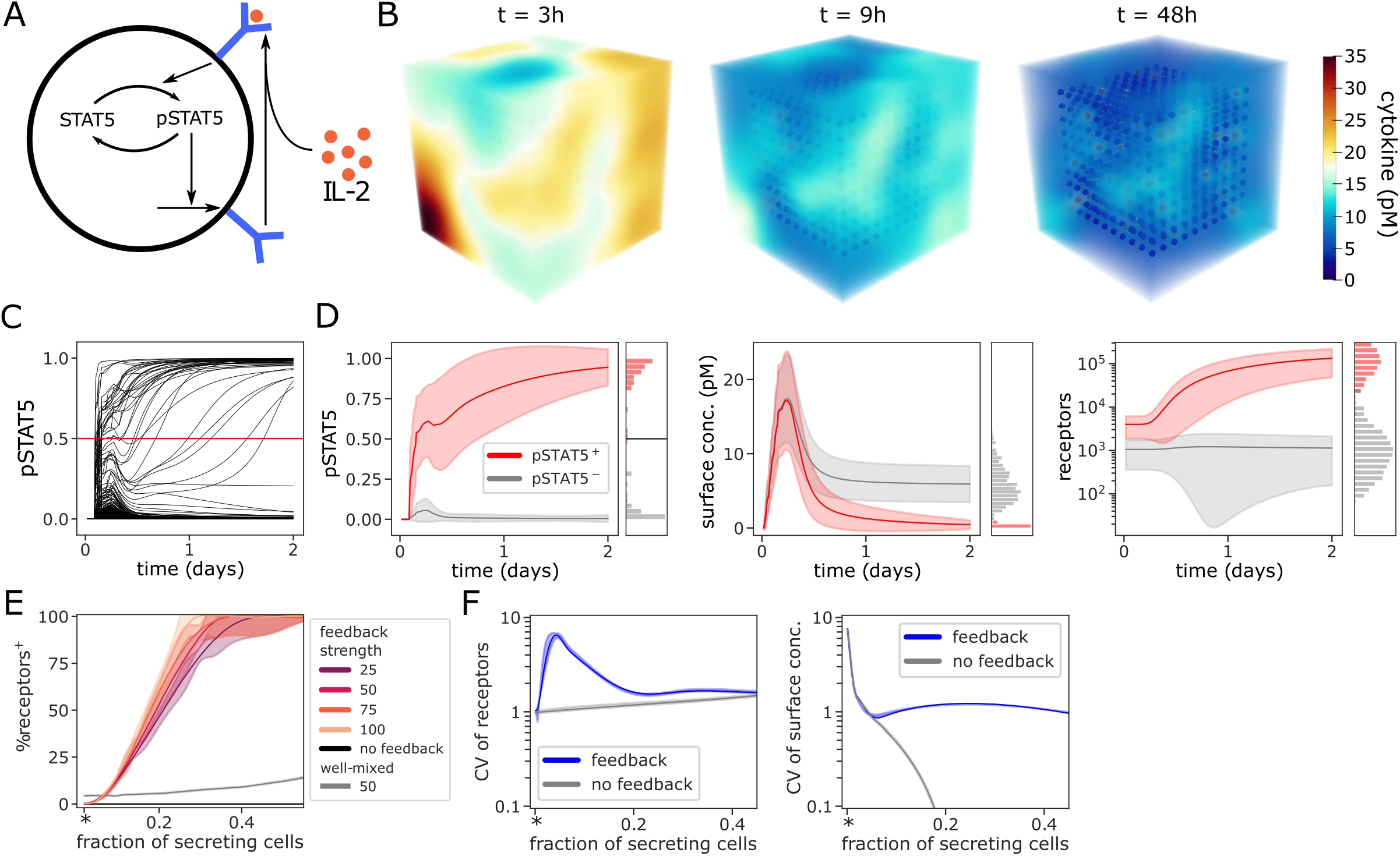
Receptor and activation kinetics in the presence of positive receptor feedback. (A) Positive feedback scheme, cytokine-receptor complexes are translated into a pSTAT5 signal that induces increased receptor expression. (B-D) Kinetic simulation of the model illustrated in panel A, shown are the resulting concentration field at indicated time points (B), the pSTAT5 signal at individual responder cells (C), and the cell-state parameters averaged separately across all activated (“pSTAT5+”) or non-activated (“+STAT5-”) cells (D), where activation is defined as pSTAT5 value above threshold (vertical line in panel C) by the end of the simulation. Shaded regions indicate SEM. (E) Parameter scan over the fraction of T_sec_ and the feedback fold change (feedback strength) for the RD-system and well-mixed model, indicating the effect on paracrine signaling effectivity by evaluation of receptor upregulation. The star indicates exactly one T_sec_. Receptor upregulation is defined by exceeding double the initial value. (F) Analysis of the effect of T_sec_ on the CV of receptors and cytokine surface concentration. Shaded regions indicate SEM over 10 simulations.

As expected based on the static simulations in Figure 2, spatial inhomogeneities were required for an effective paracrine signal and subsequent feedback-driven up-regulation of IL-2 receptor expression on responder cells (Figure 3E). A system-wide upregulation is achieved at ~25% T_sec_ already at moderate feedback strength. In contrast, the well-mixed model shows much lower activation in terms of STAT5 phosphorylation, regardless of feedback strength and even for high fractions of T_sec_ cells. Indeed, the considered positive feedback-driven dynamics induced an increase in spatial inhomogeneity in terms of both receptor expression and cytokine concentration at the surface of responder cells (Figure 3F).

### Co-localization of T_reg_ and T_sec_ cells in the vicinity of antigen-presenting cells

Recent experimental studies suggest that T_reg_ cells accumulate around clusters of cytokine secreting cells, since both cell type are stimulated by the same antigen-presenting cells (APC). T_reg_ cells differ from non-T_reg_ responder cells by higher receptor expression and cytokine uptake capacities. Therefore, co-clustering of T_reg_ cells with T_sec_ cells potentially limits activation in a fine-tuned manner, depending on the degree of localized clustering (6,9). Here, we simulated this situation by considering randomly distributed cellular positions together with a clustering algorithm that allows for a continuous scan over the clustering strength φ (Figure 4A). We found that the clustering strength in our algorithm correlates well with the silhouette score (Figure 4B), a commonly used metric quantifying association to clusters.

**Figure 4:**
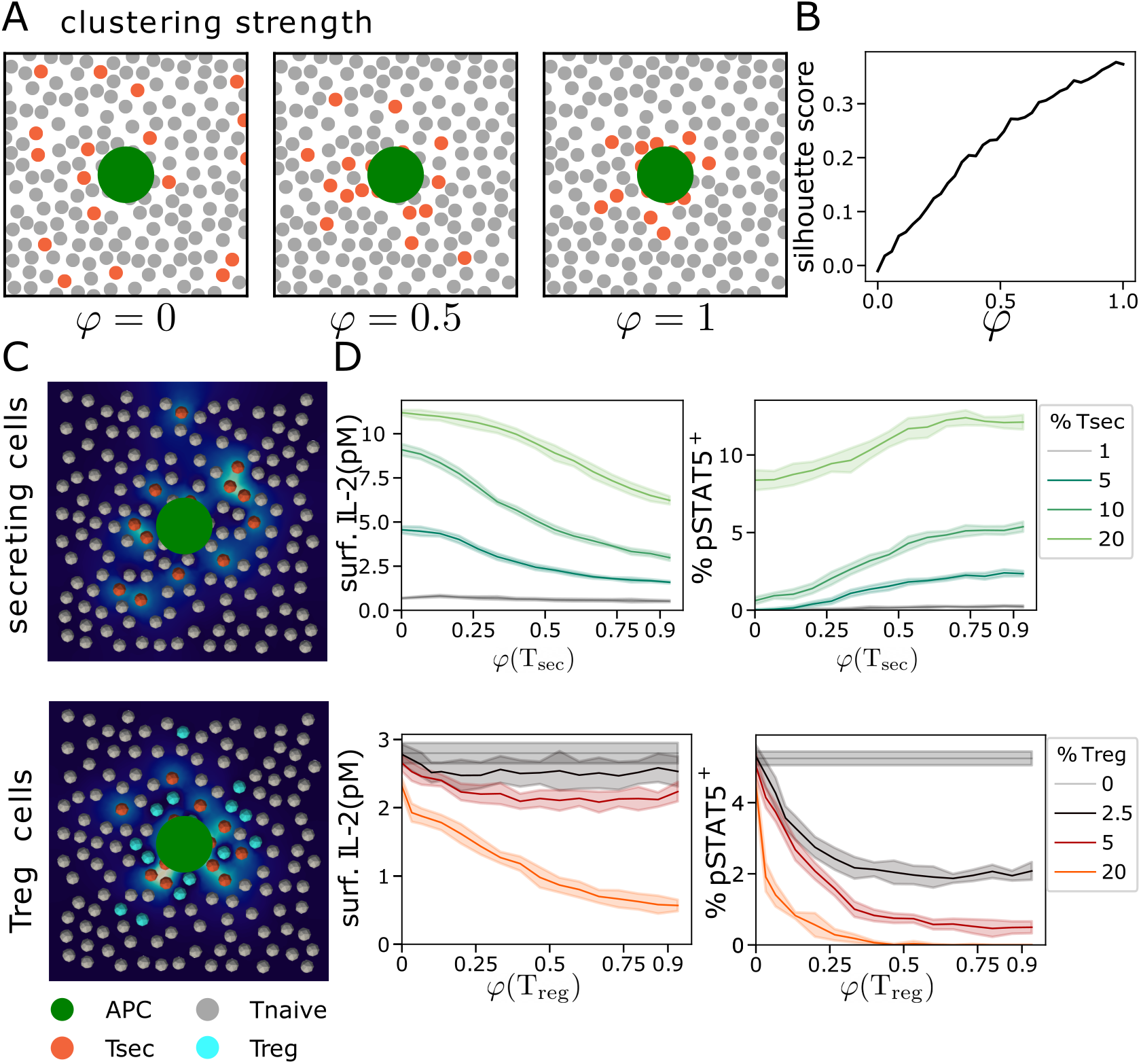
Model simulation of Treg and Tsec cell co-localization. (A) Illustration of the clustered and randomized cell layout in a two-dimensional arrangement, where one cell type (orange) is clustered around a central APC (green) with varying degrees of clustering strength φ. (B) The silhouette score is a monotonously increasing function of the clustering strength φ. (C) Illustration of T_sec_ cell clustering around a single APC at φ = 0.5 (top), and of T_reg_/T_sec_ cell clustering with φ = 1 for T_sec_ and φ = 0.5 for T_reg_ cells (bottom). (D) Effect of T_sec_ and T_reg_ clustering in the scenarios described in C. Shown are IL-2 surface concentration and paracrine signaling strength (%pSTAT5+) on responder cells at the end of simulation runs (t=100h). Shaded regions indicate SEM over 10 simulations.

We analyzed the effects of either T_sec_ clustering alone, or of T_reg_ cell clustering in the presence of highly clustered T_sec_ cells (Figure 4C). As expected, clustering of T_sec_ cells leads to reduced IL-2 concentrations and increased paracrine signaling strength (Figure 4D, top). Such effective cell-cell signaling can be mitigated by placing T_reg_ cells into the system. We found that the effect is quite limited if T_reg_ cells are placed randomly (φ = 0), even at large numbers of T_reg_ cells (Figure 4D, bottom). At least partial co-localization of T_reg_ with T_sec_ cells seems required for substantial interference with the paracrine signal, and can induce almost complete inhibition already at a fraction of 5% T_reg_ cells in our simulations.

Hence, the spatial composition of cell types with specific properties can provide another layer of fine-tuned control over the signaling range of cytokines, in addition to the amount of cytokine secreting cells, the distribution of cytokine receptor expression and cytokine-induced feedback regulation.

## Discussion

Our model combines a description of signal processing on the cellular level with a biophysical description of cytokine diffusion. Our finite-element based simulation setup allows for a systematic and scalable analysis of cytokine fields resulting from the collective information processing in T helper cell populations with varying and dynamic degrees of local collectivity (15,16). We found that the resulting spatial cytokine gradients can induce significant bifurcations in activation pattern of the naive T cell. Based on the IL-2/IL-2R system, we have demonstrated that a signaling network with positive feedback topology can further amplify the strength of spatial cytokine gradients.

Our approach, which integrates the cytokine field as a whole and does not rely on the independence of cytokine sources, is well suited to model the effects of cell clusters, where the non-linear saturation of cytokine uptake might play an important role in determining the signaling length scale. We investigated the effect and functional implications of co-localization between IL-2^+^ T_h_ - and T_reg_-cells. This phenomenon was discovered elsewhere and suggests a mechanism by which regulatory T-cells would co-localize around APCs, which present autoantigens, through TCR engagement. Our model simulations show that the spatial clustering of IL-2 secreting cells can boost the overall pSTAT5 response in responder cells, as the cytokine secretion of multiple cells combines to act on individual T cells. This effect can be counteracted by introduction of regulatory T cells into the same cell clusters, whereas uniformly distributed T_reg_ cells have little to no effect.

Overall, the spatial relationships among cytokine secreting cells and cells expressing various amounts of cytokine receptor can critically regulate the early T_h_ cell activation dynamics. Therefore, we expect that detailed quantitative analysis of the tissue architecture in immune cell compartments by both advanced imaging technologies and data-driven modeling studies will be essential for improved understanding of immune homeostasis and inflammatory pathologies (17–20).

## Methods

### Software and statistics

All model simulations were performed in Python, utilising the FEniCS library (15,16) to solve the boundary value problem on a mesh generated by Gmesh (21) in case of the spatial models, and a SciPy (22) ODE solver for our ODE model.

### Spatial continuum diffusion problem

Due to the vastly different timescales between cytokine diffusion and cell kinetics, we assume the cytokine concentration to be in steady state following the reaction-diffusion equation

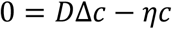

for the spatial diffusion field *c*, where D is the diffusion constant and the decay constant *η* accounts for homogeneous decay. For specific parameter values see Table 1. The design of the boundary value problem roughly follows previous spatial simulations (5) coupled to a response-time representation of cell-state dynamics (11), details will be published elsewhere.

### Well-mixed model

We assume a homogeneous extracellular volume which allows us to work with ODEs of the form

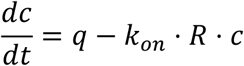

at the surface of each cell in the simulated region, where again parameters are defined as in Ref. (5). In the simulations with saturated uptake (see Section Results), the linear uptake term *k*_*on*_ ⋅ *R* ⋅ *c* is replaced by a Hill-type equation.

